# Genome-wide associations reveal human-mouse genetic convergence and modifiers of myogenesis, *CPNE1* and *STC2*

**DOI:** 10.1101/370312

**Authors:** Ana I. Hernandez Cordero, Natalia M. Gonzales, Clarissa C. Parker, Greta Sokoloff, David J. Vandenbergh, Riyan Cheng, Mark Abney, Andrew Skol, Alex Douglas, Abraham A. Palmer, Jennifer S. Gregory, Arimantas Lionikas

## Abstract

Muscle bulk in adult healthy humans is highly variable even after accounting for height, age and sex. Low muscle mass, due to fewer and/or smaller constituent muscle fibers, would exacerbate the impact of muscle loss occurring in aging or disease. Genetic variability substantially influences muscle mass differences, but causative genes remain largely unknown. In a genome-wide association study (GWAS) on appendicular lean mass (ALM) in a population of 85,750 middle-age (38-49 years) individuals from the UK Biobank (UKB) we found 182 loci associated with ALM (*P*<5×10^−8^). We replicated associations for 78% of these loci (*P*<5×10^−8^) with ALM in a population of 181,862 elderly (60-74 years) individuals from UKB. We also conducted a GWAS on hindlimb skeletal muscle mass of 1,867 mice from an advanced intercross between two inbred strains (LG/J and SM/J) which identified 23 quantitative trait loci. 38 positional candidates distributed across 5 loci overlapped between the two species. *In vitro* studies of positional candidates confirmed *CPNE1* and *STC2* as modifiers of myogenesis. Collectively, these findings shed light on the genetics of muscle mass variability in humans and identify targets for the development of interventions for treatment of muscle loss. The overlapping results between humans and the mouse model GWAS point to shared genetic mechanisms across species.

## Introduction

Skeletal muscle plays key roles in locomotion, respiration, thermoregulation, maintenance of glucose homeostasis and protection of bones and viscera. The loss of muscle due to aging, known as sarcopenia, affects mobility and can lead to frailty and deterioration of quality of life^1^. The risk of disability is 1.5 to 4.6 times higher in the sarcopenic elderly than in the age-matched individuals with normal muscle mass^2^. However, lean mass, a non-invasive proxy for muscle mass, differs by more than two-fold between healthy adult individuals of same sex, age and height^3^. Therefore, we hypothesize that differential accretion of muscle mass by adulthood may influence the risk of sarcopenia and frailty later in life.

Genetic factors contribute substantially to the variability in lean mass in humans, with heritability estimates of 40 – 80 %^4^. A continuous distribution of the trait and data obtained from animal models^5–7^ indicates a polygenic causality. However, thus far, genome-wide association studies (GWAS) have implicated fewer than a dozen genes, explaining a small fraction of this heritability^8; 9^. Limited sample size in early studies^10–14^ and the effects of confounders such as subject age^8^, size of the skeleton and lean mass components (non-fat organs and tissues, heterogeneity of muscle fibers) have hindered detection of genes. The UK Biobank is a resource of demographic, phenotypic and genotypic data collected on ∼500,000 individuals^15^. It includes the arm and leg lean mass, body composition and morphometric information, providing a model for improving our understanding of the genetic basis for variability in muscle mass. Skeletal muscle mass, however, changes over the course of an individual’s lifespan. It reaches a peak around mid-twenties and remains largely stable through mid-forties, before succumbing to gradual decline, which accelerates after about 70 years of age^16^. There is a substantial degree of individual variability in the slope of muscle change across both the increasing and decreasing phases of the lifespan trajectory^17^. Both the trajectory itself and the slope of individual variability may impede identification of genes.

The indirect estimates of lean mass impose limitations because muscle mass is not an exclusive contributor to this variable. Furthermore, the cellular basis of variability in muscle mass (i.e. if it is caused by the differences in the number of constituent muscle fibers, their size, or both) remains poorly understood. Using the laboratory mouse circumvents a number of those limitations. The mouse shares approximately 90% of the genome with humans^18^, and dissection permits analyses of traits that are difficult to study directly in humans, such as the mass of individual muscles ^6; 7; 19^ and whole-muscle fiber characteristics^20; 21^. The phenotypic differences between the LG/J and SM/J mouse strains make them particularly attractive for complex trait analyses^22–24^. LG/J mice were selected for large body size^25^, while SM/J mice were selected for small body size^26^. The second filial generation (F2) of intercross derived from the LG/J and SM/J strains (LGSM)^6; 27^ and an advanced intercross line (AIL) of the LGSM (LGSM AIL), developed using a breeding strategy proposed by Darvasi and Soller^28^, led to multiple quantitative trait loci (QTLs) for hindlimb muscle mass^6; 27^. However, these QTLs still encompass tens or even hundreds of genes and require further prioritizing. We hypothesized that the detection power of a modest sample size of the LGSM AIL and the superior resolution of a human cohort will facilitate identification of the quantitative trait genes (QTGs) underlying muscle QTLs.

The aim of this study was to identify the genomic loci and the underlying genes for variability in skeletal muscle mass and to assess their effects in the elderly. We addressed this in three stages: (1) we conducted a GWAS in a human cohort of middle-aged individuals from the UK Biobank and tested the effect of the identified set of loci in an elderly cohort; (2) we conducted a GWAS on hindlimb muscle mass in a population of LGSM AIL mice. (3) In the final stage, we nominated candidate genes by comparing mouse and human loci and validated the myogenic role of selected candidates *in vitro*.

## Methods

### Stage one: Genome mapping in human populations

#### UK Biobank cohort

The population in this study consisted of 316,589 adult individuals of 37 to 74 years of age (project ID: 26746). We drew this cohort from the UK Biobank (UKB) project^15^; all participants recruited were identified from the UK National Health Service (NHS) records and attended a baseline visit assessment between 2006 and 2010. During the assessment, participants gave written consent, answered a questionnaire, and were interviewed about their health and lifestyle. Blood samples and anthropometric measurements were collected from each participant. Assessments were conducted at 22 facilities in Scotland, England, and Wales.

We divided the sample into middle-aged and elderly cohorts. The middle-aged cohort consisted of 99,065 adults ranging from 38 to 49 years of age; based on previous studies we assumed that these individuals were not affected by sarcopenia^29^. We excluded 3,520 participants that were reported to be ill with cancer, pregnant, or had undergone a leg amputation procedure, as well as individuals with discordant genetic sex and self-reported sex records. In addition, we excluded non-white Europeans (self-reported) from the analyses (n = 9,599) and individuals without imputed genotypes. We retained a total of 85,750 adult individuals (46,353 females and 39,397 males) for further analyses.

The elderly cohort consisted of 217,524 adults ranging from 60 to 74 years of age. We selected this cohort to test if the effect of the genetic variants identified on middle-aged individuals could also influenced phenotypes later in life. We excluded 35,662 individuals based on the same criteria used for the middle-aged cohort. After exclusions, the elderly cohort included 181,862 individuals of 60 to 73 years of age (94,229 females and 87,633 males; Table S1).

#### UK biobank traits

We used the data for standing height (UKB field ID: 50), sitting height (UKB field ID: 20015), whole body fat (UKB field ID: 23100), arm lean mass (UKB field ID: 23121 and 23125), and leg lean mass (UKB field ID: 23113 and 23117) measured as part of the UK Biobank project. Body composition measurements were taken using bioelectric impedance (this was preferred to the DXA scan data because of the substantially larger number of phenotyped individuals). We calculated leg length by subtracting sitting height from standing height (all measurements were recorded in cm). Because lean mass in the limbs primarily consists of skeletal muscle tissue, we used ALM as a proxy for muscle mass. We calculated ALM as the sum of the muscle mass of two arms and two legs. We checked that all traits were normally distributed by examining the QQ-plot and histogram of residuals from a simple linear model that included sex as a covariate. Residuals were normally distributed, and we did not transform any of the traits.

#### UK biobank genotypes

We obtained genotype data for all participants from the UKB v3 genotypes release^30^, which includes genotype calls from the Affymetrix UK BiLEVE Axiom array and the Affymetrix UK Biobank Axiom array, and imputed genotypes from the UK10K and 1000 Genomes Phase 3 reference panels^31^. We kept all imputed genotype data (21,375,087 genetic variants (SNPs, Indels and structural variants)) with MAF > 0.001 and imputation quality > 0.30. The software (BOLT-LMM v2.3.4) ^32^ we used to perform GWAS was developed for large data sets (i.e.: UK Biobank cohort) and it was only tested for human cohorts, which have different LD patterns from animals; BOLT-LMM uses a linear mixed model, which have been shown to successfully control for confounding due to population structure or cryptic relatedness in individuals (related and unrelated) from the UK biobank^33–36^. For these reasons, we used BOLT-LMM v2.3.4 for the analyses of human data only.

#### Appendicular lean mass GWAS

We used BOLT-LMM (v2.3.4)^37^ to perform a GWAS for ALM in the middle-aged cohort. The linear mixed model (LMM) approach implemented in BOLT-LMM is capable of analyzing large data sets while also accounting for cryptic relatedness between individuals. Specifically, BOLT-LMM calibrates the association statistics using a linkage disequilibrium (LD) score regression approach^38^; this allowed us to evaluate the impact of confounding factors on the GWAS test statistics^38^ and calibrate them accordingly. In the absence of confounding factors, *P* values should not be inflated, and the LD score regression intercept should be equal to 1^38^. The LD Score regression intercept in this study was 1.051 ± 0.007, suggesting minimal inflation of *P* values due to linkage between markers. After calibrating the test statistics, the mean χ^2^ of the ALM GWAS was 1.29 and lambda (λGC) or genomic control inflation factor was 1.20 (Figure S1), which indicated polygenicity of the trait as described by Bulik-Sullivan and colleagues^38^.

We also assessed population structure by running principal component analysis on the genotype calls. We included sex, leg length, whole body fat, and the first four principal components as fixed effects in the LMM used for the ALM GWAS. Sex was included to account for differences in muscle mass caused by higher testosterone levels in males^39^. Testosterone is a potent stimulator of muscle growth and if systematically varied in males, it can also influence muscle mass (e.g. as a result of hypogonadism^40^). However, if there was a common genetic basis for such variability it could be captured in the association analysis. It needs to be noted that inclusion of sex as a covariate would not permit capturing sex-by-locus interactions. Identification of sex specific loci, albeit of interest, was not attempted due to complexity posed by the number of genetic markers and the sample size. An outcome of a GWAS would also depend of the complexity of mechanisms affecting the phenotype and adjustments included in a model^8; 41^. Leg length and whole body fat were included because they are biologically related to muscle mass: longer bones result in longer muscles, while fat shares part of its developmental origin with skeletal muscle tissue^42^. Furthermore, each of these traits is correlated with muscle mass. An association was considered statistically significant if its *P* < 5 × 10^−8^ (α = 0.05). This threshold is the standard for GWAS of complex traits^43; 44^.

We obtained variance components and SNP heritability estimates of ALM from the middle-aged cohort using BOLT-REML^37^. The BOLT-REML method robustly estimates the variance of genotyped SNPs and fixed effects on the LMM. As described by Loh et al. ^45^, BOLT-REML partitions the SNP heritability across common alleles; hence, the additive variance is calculated as the cumulative variance of genotyped SNPs.

#### Phenotypic variance explained by ALM loci

We defined ALM genomic loci using the web-based platform Functional Mapping and Annotation of Genome-Wide Association Studies (FUMA GWAS^46^). A key feature of this tool is the identification of genomic regions and independent genomic signals based on the provided summary statistics of a GWAS depending on LD structure; this process is automated using pairwise LD of SNPs in the reference panel (1000 genomes project phase 3 EUR ^47^) previously calculated by PLINK^48^. We provided to FUMA GWAS the summary statistic of our GWAS on ALM with the following parameters: 250kb window (maximum distance between LD blocks), r^2^ > 0.6 (minimum r^2^ for determining LD with independent genome-wide significant SNPs used to determine the limits of significant genomic loci), MAF > 0.001 (minimum minor allele frequency to be included in the annotation), *P* < 5 × 10^−8^ (threshold of significantly associated variants). We refer to the identified regions and the independent signals as loci throughout the text.

We used the top variant (based on the outcome from FUMA^46^) of each locus identified to estimate the proportion of phenotypic variance explained by each locus. We estimated phenotype residuals using a model that included the fixed effects and principal components described above. We then regressed the residuals on the genotype of the top SNP in a linear model. We estimated the coefficients of determination and reported them as the proportion of phenotypic variance explained by each locus.

#### Genetic effects in the elderly cohort

We tested the combined effect of all 182 genome-wide significant ALM loci identified in the middle-aged cohort in the elderly cohort using the top SNP at each locus. We used PLINK2^48^ to extract genotype dosages for each variant identified in the middle-aged GWAS in the elderly cohort. We then estimated a ‘genetic lean mass score’ for each individual using the following procedure. First, we estimated the contribution of each variant on the phenotype as a product of the SNP effect size obtained from BOLT-LMM (β, calculated based on the reference allele) and the genotype dosage. Second, we calculated the ‘lean mass score’ for each individual as the sum of the products for all selected variants. We ranked the resulting distribution of lean mass scores in ascending order and partitioned it into five quantiles. We used ALM without any adjustment (raw ALM) because estimates of effects size already accounted for sex, whole body fat and leg length differences. However, since the raw ALM did not meet the assumption of normality, we used a Kruskal-Wallis test (non-parametrical) to evaluate the difference in the median of the phenotypes between the quantiles, and a Wilcoxon test (non-parametrical) for pairwise comparisons between quantiles. We conducted five replicates of a negative control test that consisted of randomly selecting a subset (n ∼ 185) of non-significant SNPs in the middle-aged cohort and generating ‘lean mass score’ as described above for the elderly cohort; this set of SNPs had MAF > 0.001.

We also aimed to replicate the individual variants effects on the ALM of the elderly cohort. We checked normality of ALM in the elderly cohort as described for the middle-aged cohort. We tested a subset of genetic variants (n=17,914,406) selected based on their MAF > 0.001 and imputation quality > 0.3, and we used the same LMM, fixed covariates, and genome-wide significance threshold (*P* < 5 × 10^−8^) as described for the middle-aged cohort. We conducted a Fisher’s exact test to evaluate if overlapping loci between the middle-aged and elderly cohorts were significantly different from random. The null hypothesis was rejected at P < 0.05 (two-tailed).

#### Genomic regions tagged by loci

We used the ‘biomaRT’ package in R^49; 50^ to retrieve gene and regulatory element annotations at the genomic position of each statistically significant SNP (*P <* 5 × 10^−8^) and Polyphen 2^51^ and SIFT^52; 53^ to predict the functional consequences of each SNP. We retrieved additional information about the positional candidate genes and their expression levels from Ensembl^54^ (release 94 - October 2018) and the Genotype Tissue Expression Project (GTEx) portal^55^ (See Web Resources).

### Stage two: LGSM AIL mouse cohort and GWAS

To maximize QTL detection power, we combined three cohorts of LGSM AIL mice for the second stage of this study (n = 1,867). The LGSM AIL was initiated by Dr. James Cheverud at Washington University in St. Louis ^56^. Cohort 1 included 490 mice (253 males and 237 females) from LGSM filial generation 54 (F34). Phenotype data was collected from these mice between 80-102 days of age. Cohort 2 consisted of 506 male mice (∼ 84 days of age) from filial generation 50-54 (F50-54). Cohort 3 includes 887 mice (447 males and 440 females) from filial generation 50-56 (F50-56); between 64 - 111 days of age. Mice were housed at room temperature (70 - 72°F) at 12:12 h light-dark cycle, with 1-4 same-sex animals per cage and with *ad libitum* access to standard lab chow and water.

#### Mouse traits and genotypes

We collected muscle phenotypes after the animals were sacrificed and frozen. We dissected four muscles and one long bone (tibia or femur) from each mouse at the Pennsylvania State University (n = 584) and the University of Aberdeen (n = 1,283). Each of the four muscles exhibits a different proportion of muscle fiber types and often revealed muscle specific QTLs^6; 7; 19; 27^.The dissection procedure involved defrosting the carcasses and removing the muscles (TA, EDL, gastrocnemius and soleus) and tibia from the hind limbs under a dissection microscope. We weighed the muscles to 0.1-mg precision on a Pioneer balance (Pioneer, Ohaus) and measured long bone length of the hindlimb (mm) using an electronic digital calliper (Powerfix, Profi). We quantile normalized all LGSM AIL traits before mapping QTLs.

Cohort 1 was genotyped using a custom SNP genotyping array^57^. These SNPs (n=2,965) were evenly distributed along the autosomes (*Mus musculus* genome assembly MGSCv36 (mm8)). The median distance between adjacent SNPs was 446 Kb, and the maximum was 18 Mb. Large gaps are due to regions identical by descent between the LG/J and SM/J founders^58^. Cohort 2 was genotyped at 75,746 SNPs (73,301 on the autosomes and 2,386 on X and Y) using the MEGA Mouse Universal Genotyping Array (MegaMUGA; *Mus musculus*) genome assembly MGSCv37 (mm9)); after removing SNP markers that are not polymorphic between the LG/J and SM/J strains we retained 7,168 autosomal SNPs for subsequent analyses. The median distance between adjacent SNPs was 126.9 Kb and the maximum distance was 15 Mb for all chromosomes except for chromosomes 8, 10, and 14, which had distances of 19, 16, and 16 Mb, respectively. We used a conversion tool in Ensembl to convert SNP positions from mm8 and mm9 to *Mus musculus* genome assembly GRCm38 (mm10). Cohort 3 genotypes were obtained from Gonzales and colleagues^59^. These genotypes were generated using genotyping by sequencing. This approach has been recently used and described in detail^19^. Only autosomal SNPs known to be polymorphic in the LG/J and SM/J founder strains (n=523,027; mm10, build 38) were retained for subsequent analyses. We combined the genotype data from Cohorts 1-3 using PLINK v.1.9 and imputed missing genotypes using BEAGLE v.4.1^60^. For these steps, we used a reference panel obtained from whole genome sequencing data of the LG/J and SM/J strains^58^. Dosage estimates (expected allele counts) were extracted from the output and used for the GWAS; these estimates captured the degree of uncertainty from the imputation procedure. To ensure the quality of the genotype data, we excluded SNP genotypes with MAF < 0.20 (because it is an AIL, almost all SNPs have MAF > 0.2) and dosage R^2^ < 0.70 (dosage R^2^ corresponds the estimated squared correlation between the allele dosage and the “true allele dosage” from the genetic marker, and is used as a measure of imputation quality). After applying these filters, we retained 434,249 SNPs.

#### Mouse association analyses

Population structure can potentially lead to a rise of false positive associations^61; 62^. The LMM approach is used to map QTLs while dealing with confounding effects due to relatedness^57; 63; 64^. We used the LMM method implemented in the software GEMMA (genome-wide efficient mixed-model association)^65^ to analyze the mouse phenotypes. In our LMM model we included the genotypes, a set of fixed effects described later in this section, and a polygenic effect to deal with population structure.

The polygenic effect is a random vector which was derived from a multivariate normal distribution with mean zero and a n × n covariance matrix σ^2^λK; where n is the number of samples. The relatedness matrix K was defined by the genotypes. The two parameters, σ^2^ and λ, were estimated from the data by GEMMA; they represent the polygenic and residual variance components of the phenotypic variance, respectively.

#### Relatedness matrix and proximal contamination

We used the genotype data to estimate the relatedness matrix K, which was part of the covariance matrix. Although genotype-based and pedigree-based K matrices yield very similar results^66; 67^, we have shown that in general, genotype-based estimates are more accurate^66; 68–70^. We constructed the relatedness matrix as K = XX^′^/p, where X is the genotype matrix of entries x_ij_and n × p dimensions, p is the number of SNPs.

The relatedness matrix K was estimated taking into account the potential problem of proximal contamination^67^, which involves loss of power due to including genetic markers in multiple components of the LMM equation. Furthermore, because of LD, markers in close proximity to the genetic marker that is being tested can also lead to deflation of the *P* values^59; 68^. To avoid this problem, the K matrix was estimated by excluding from the calculations the SNPs within the chromosome that was analyzed (this approach is termed leave one chromosome out (LOCO)), therefore, K matrix was slightly different for each chromosome.

#### Genetic and fixed effects

We did not include non-additive effects in the LMMs used for GWAS in the LGSM AIL. Our previous studies^6^ suggest that musculoskeletal traits in this population are mostly influenced by additive loci, and by ignoring dominance effects we avoid introducing an additional degree of freedom, hence potentially avoiding a decrease of power to detect QTLs.

To analyze the muscle weights of the combined data, we used four fixed effects in the LMM: sex, dissector of the samples, age, and long bone length of the hindlimb. We selected these variables after using a linear model to estimate their effect on the four muscles; only statistically significant effects were included (P < 0.01). Sex and dissector were included as binary variables; whereas age and long bone were included as continuous variables. Including long bone length of the hindlimb allowed us to capture genetic effects associated with variation in muscle weight *per se* (as opposed to genetic effects on bone length)^19^. In other words, failing to include long bone as a covariate would yield QTLs that are more likely to be genetic contributors to general growth of the skeleton instead of specifically muscle. We used two bones for the long bone variable, for cohort 1 femur, and for cohorts 2 and 3 tibia. Based on personal communication with Dr. Cheverud, the femur and tibia bones were found to be positively and highly correlated (*r* > 0.8) in LGSM AIL (F34). We did not include generation (*r* = 1) and bone type of each cohort (*r* = 1) as fixed effects since the dissector variable functioned as a proxy for these two variables. Body weight was not used as a fixed effect because muscle weight accounts for a considerable amount of the body weight.

#### SNP heritability

To estimate the SNP heritability or proportion of phenotypic variance explained by all genotypes, we used the n × n realized relatedness matrix K, which was constructed using all the available genotypes. We extracted the SNP heritability from the QTL mapping outputs of the LGSM AIL cohort described before; GEMMA provides an estimate of the heritability and its standard error^65^. The SNPs available to estimate the heritability do not capture all genetic causal variants, hence the SNP heritability underestimates the true narrow sense heritability.

#### Threshold of significance and QTLs intervals

The *P* values estimated from the likelihood ratio test statistic performed by GEMMA were transformed to –log_10_ *P* values. We calculated a threshold to evaluate whether or not a given SNP significantly contributes to a QTL. We estimated the distribution of minimum *P* values under the null hypothesis and selected the threshold of significance to be 100(1 – α)^th^ percentile of this distribution, with α = 0.05. In order to estimate this distribution, we randomly permuted phenotypes 1,000 times, as described previously ^6; 7; 19; 71^. We did not include the relatedness matrix in the permutation tests due to computational restrictions, and because, past studies have found that relatedness does not have a major effect on the permutation test^6; 7^.

We estimated QTL intervals in three steps. 1) We used Manhattan plots to identify the top SNP within each statistically significant region (SNP with highest –log_10_ *P* values), which we refer to as the peak QTL position. 2) We transformed *P* values from each analysis to LOD scores (base-10 logarithm of the likelihood ratio). 3) We applied the LOD interval function implemented in the r/qtl package^72^ to the regions tagged by each peak SNP, and obtained the QTL start and end positions based on the 1.5 LOD score interval. 1.5 LOD intervals are commonly used to approximate the ∼ 95% confidence interval of mouse QTLs^5; 73^. The 1.5 LOD interval estimation is comparable to the 95% CI in the case of a dense marker map^74^; hence, its coverage depends on the location of the peak QTL marker relative to the adjacent genotyped markers. We estimated the direction of the QTL effect by calculating the phenotypic mean of each allele based on the peak SNP of each QTL. We adjusted the phenotypic means and standard errors by fitting the fixed effects used in the association analyses in a linear model.

We explored the QTL intervals to identify genes that potentially affect hindlimb muscle mass. We retrieved the genomic location of all genes located within the intervals using the BioMart database through the ‘biomaRT’ package implemented in R^49; 50^.

#### Meta-analysis in the LGSM AIL mice

Although we adjusted our GWAS analyses on the LGSM AIL mice for confounding effects, it was possible that uncontrolled factors could have affected the phenotypes. Therefore, we conducted an additional meta-analysis on the three LGSM AIL cohorts. We first analyzed each cohort separately using the same approach as for the combined data, except the dissector variable was not used as a covariate because it was largely confounded with the cohort. We extracted *P* values, estimated SNP effects and standard errors at each scanning locus. We considered two popular meta-analysis approaches: the inverse variance-weighted average and the weighted sum of z-scores^75–77^. For weighted sum of z-scores, we tried two weighing schemes, i.e., the sample size and squared root of the sample size, and found the results were very similar, and were slightly better than that of the inverse variance-weighted average. Therefore, we chose to report the result of the weighted sum of z-scores with the squared root of the sample size being the weight as suggested in meta-analysis literature^78^. The test statistic (*Z*) for each SNP was constructed as follows:

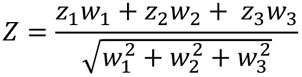

where *z_i_* is the z-score that was obtained by transforming the likelihood ratio test *P* value and *w_i_* is the squared root of the sample size in cohort *i*=1 to 3. We compared this result with the combined GWAS of the LGSM AIL. This statistical analysis was performed in R^79^.

#### Overlap of mouse and human results

The significantly associated muscle QTLs (mice) and lean mass loci (humans) were compared by exploring the genomic regions and genes tagged in each analysis. We used a Fisher’s exact test to evaluate whether the number of overlapping loci from the human and mouse analyses exceeded the number expected by chance; the null hypothesis was rejected at P < 0.05.

#### Gene validation using siRNA in C2C12 myoblasts

To validate efficiency of siRNA-mediated gene knockdown, the C2C12 cells were lysed and RNA isolated using RNeasy mini kit (QIAGEN) following manufacturers recommendations. Concentration was assessed using NanoDrop (Thermo Scientific) spectrophotometer and ∼1.5 µg of RNA was applied to 1.5% agarose gel to validate its integrity. The cDNA was synthesized using random primers (Invitrogen) and SuperScript II reverse transcriptase (Invitrogen). Quantitative PCR for expression of the target *Cpne1*, *Sbf2* and *Stc2* and the reference *Actb* was carried out in duplicates on LightCycler 480 II (Roche) using SYBR green Master mix (Roche), 10 ng cDNA and 0.5 µM forward and reverse primers (Table S2). Quantification of gene expression was performed using the comparative Ct method^80^.

C2C12 myoblasts, validated for differentiation, were seeded on 8-chamber slides (Lab-Tek II), batch 1, and 13 mm diameter Thermanox Plastic coverslips (Thermo Fisher Scientific), batch 2, at 100 cells/mm^2^ in high glucose growth medium (D5671, Sigma), containing 10% foetal calf serum and 2% glutamine. Next day the cells were washed with phosphate-buffered saline (PBS) and transferred to differentiation medium (D5671, Sigma) supplemented with 10 nM siRNA and Lipofectamine RNAiMAX (Invitrogen) as per manufacturer protocol. We used the following siRNAs (Life Technologies): negative control #1, s113938 and 93494 (*Cpne1*), 151885 and 151886 (*Stc2*), s115441 and s115442 (*Sbf2*). The treatment achieved expression knockdown by 55-70%. The differentiation medium with 10nM siRNA and Lipofectamine RNAiMAX were replaced once, after 3 days of incubation. After 6 days of incubation, cells were fixed in 4% paraformaldehyde (PFA). We examined 8 cultures for *Stc2* and 12 for the remaining genes (equally divided between the two siRNAs) and negative control that were generated in two batches on separate occasions.

Cells were washed in PBS, fixed in 4% PFA for 15 min, PBS washed again and permeabilized for 6 min with 0.5% Triton X-100 in PBS. The cells were then blocked for 30 min in blocking buffer (10% foetal calf serum in PBS) and incubated overnight at 4 °C with primary anti-myosin heavy chains antibody (Monoclonal Anti-Myosin skeletal, Fast, Clone My-32, Mouse Ascities Fluid, M4276, Sigma-Aldrich) diluted (1:400) in PBS. After three washes in 0.025% Tween-20 in PBS at room temperature, secondary donkey anti-mouse IgG H&L antibody (ab150109, abcam) conjugated to fluorescent dye (Alexa Fluor 488) in PBS (1:400) were applied and incubated for 90 minutes. Following three washes in 0.025% Tween-20 in PBS cells were incubated in 300 nM DAPI in PBS for 15 min. After that cells were covered by coverslip using Mowiol 4-88 (Sigma-Aldrich), sealed with nail polish, and stored at 4 °C in the dark.

Slides were scanned using Axioscan Z1 slide scanner (Zeiss) using X20 magnification. The entire 0.7 cm^2^ chamber of a slide or a coverslip was imaged using the wavelength spectrum band of 353-465 nm and 493-517 nm and exposure time 4 ms and 100 ms for DAPI and Alexa Fluor, respectively, at 50% Colibri 7 UV-free LED light source intensity. Alexa Fluor and DAPI channel images of a rectangular area free of artefacts and covering at 14-91% of a chamber of batch 1 and 70% of a coverslip of batch 2 were exported separately for analyses with Fiji^81^. Note that the rectangle area of the majority of batch 1 samples (88%), covered more than 40% of the cell culture. A sensitivity analysis testing the exclusion of small coverage images (14-31%) from the statistical analyses described below, showed results comparable to the analysis of all samples; therefore, we reported significance values (*P* values) corresponding to the statistical analysis of all samples.

Three indices characterizing the effect of treatment on myogenesis were quantified in an unbiased, automated analysis of the entire exported area: 1) percentage of fluorescent area in the Alexa Fluor channel, reflecting the level of myosin expression, and 2) the longest-shortest-path reflecting the length and number of myotubes (Figure S2). The longest-shortest-path analysis was carried out using the Analyze Skeleton plugin^82^ and the shortest path calculation function^83^ implemented in Fiji^81^. We carried out the image analyses on a Linux computer and we allocated 190 GB of RAM for these analyses. The myotube threshold was set at 103.97 µm for batch 1 and 191.63 µm for batch 2, i.e. the mean (batch 1: 54.34 µm, batch 2:100.95 µm) plus 3 standard deviations (batch 1: SD = 16.54 µm, batch 2: SD = 30.23 µm) of the length of mononucleated and myosin expressing myocytes (n=35) measured in the negative control #1 cells. The myotube length variable did not follow normality, therefore quantile normalization was applied to the variable. All statistical analyses were adjusted for the image area of each sample and batch of cells, by fitting a linear model on the three indices investigated; all subsequent statistical analyses were conducted on the residuals, which met the assumptions of normality and homoscedasticity of residuals. The effect of gene knockdown on these indices was assessed using ANOVA. After, a t-test was carried out to evaluate the mean differences between the control group and the gene knockdown groups. In addition, we evaluated the myosin expressing area (as percentage of the total) present within each knockdown versus control groups using ANOVA.

#### Data availability

The human data used for this study can be obtained upon application to the UK biobank project^15^.

## Results

### Over 180 genomic loci associated with appendicular lean mass in humans

The appendicular lean mass (ALM) ranged from 12.2 to 41.6 kg and 15.3 to 54.5 kg in healthy middle age females and males, respectively (Table 1). SNP heritability estimates indicated that 36% of phenotypic variability was due to genetic factors. The GWAS analysis results presented in Figure 1 revealed 6,693 autosomal variants (MAF > 0.001) associated (*P* < 5 × 10^−8^) with ALM (Table S3). The associated variants tagged 331 genes and 753 regulatory elements. We used the Functional Mapping and Annotation of Genome-Wide Association Studies (FUMA GWAS^46^) to define genomic regions containing the associated variants, and we identified 77 of them that were on average 0.40 Mb long and contained 182 independent signals (Table S4). We refer to the identified regions and the independent signals as “loci” throughout the text. The 182 loci identified indicate that ALM is influenced by multiple genetic elements. The LD score intercept that we estimated during this ALM GWAS (1.05 ± 0.007 (mean ± SE)) provides further evidence suggesting polygenicity. Cumulative effects of these loci explained 24% of SNP heritability.

**Figure 1.**
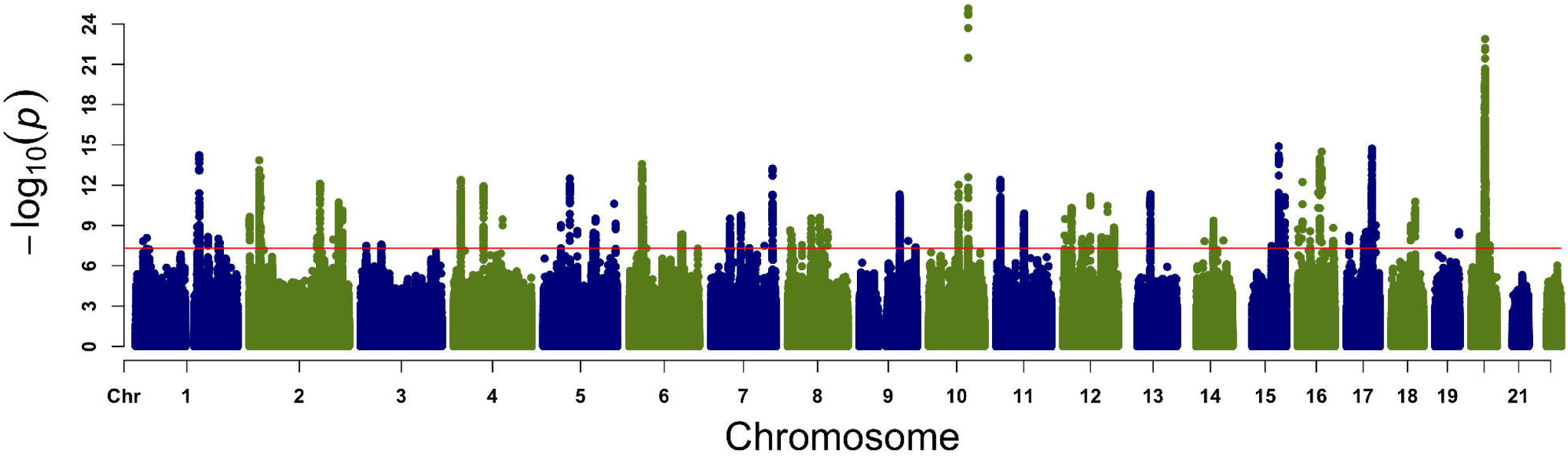
Map of genome associations with the appendicular lean mass (ALM) of humans. Genome-wide association study (GWAS) on the ALM of middle-aged adults from the UK Biobank. Significance level is presented on the vertical axis, while the chromosomal position of each genetic marker is shown on the horizontal axis. Red line across the plot represents the genome-wide threshold of significance (*P* < 5 × 10^−8^). This plot shows the association of variants with MAF > 0.001.

**Table 1.**
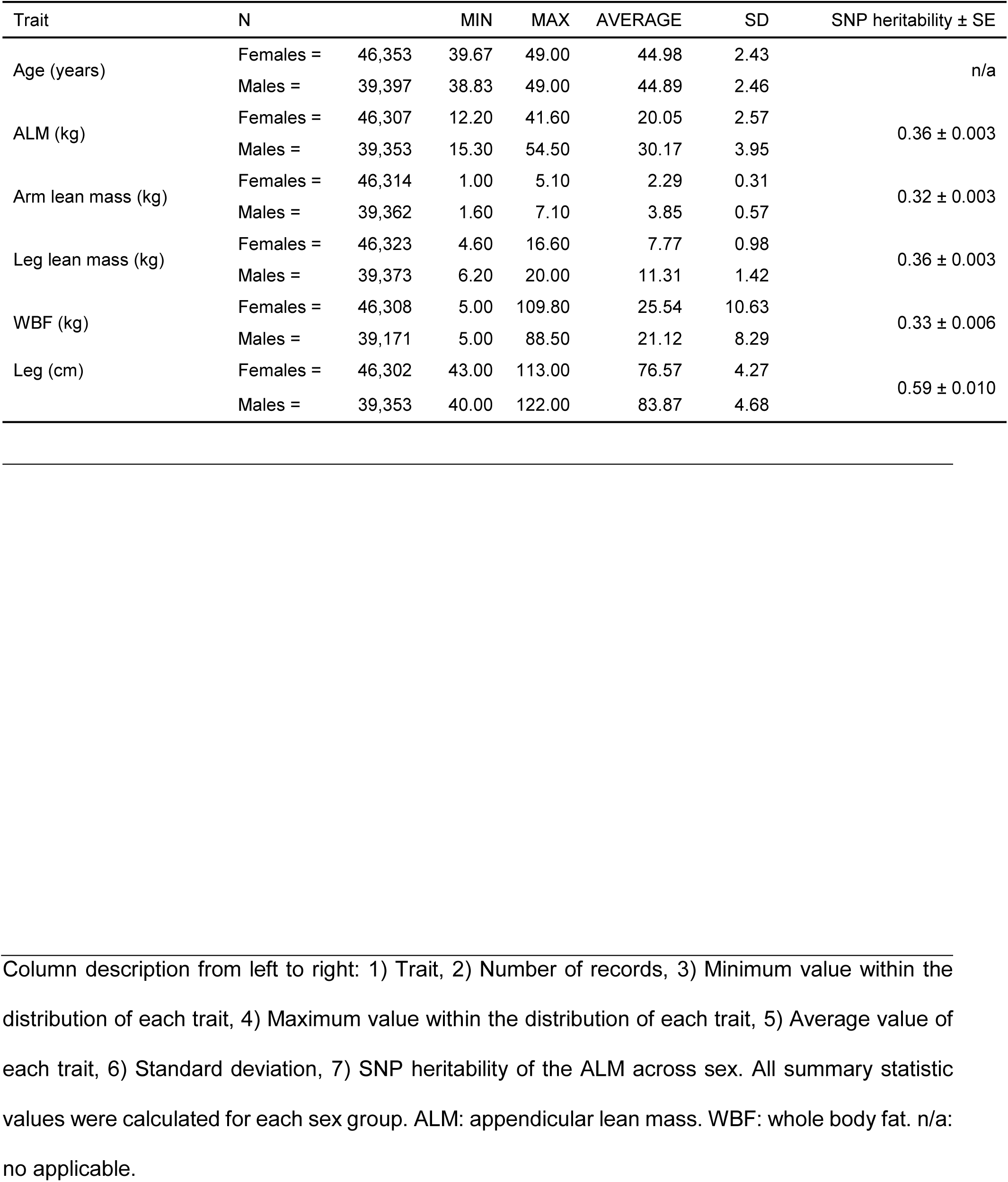
Summary of the middle-aged cohort

### 78% of the same loci affect appendicular lean mass in older adults

As expected due to the aging effect on skeletal muscle, the ALM in the cohort of elderly declined by 4 and 8% in comparison to the middle-age cohort of females and males, respectively (*P* < 2×10^−16^). A more prominent decline in males is consistent with earlier reports^84^. We then used a ‘genetic lean mass score’ (see Methods for details) to test if the identified 182 loci contributed to ALM variability in the elderly population. The genetic lean mass score had a statistically significant overall effect (χ^2^ = 376.13, df = 4, *P* = 3.99×10^−80^) on ALM variability in the elderly population (Figure 2). On average, individuals with the highest genetic lean mass score had 0.73 kg, or 3.2%, more ALM compared to those with the lowest scores (Figure 2). Negative controls showed no statistically significant effects (Table S5 and Figure S3).

**Figure 2.**
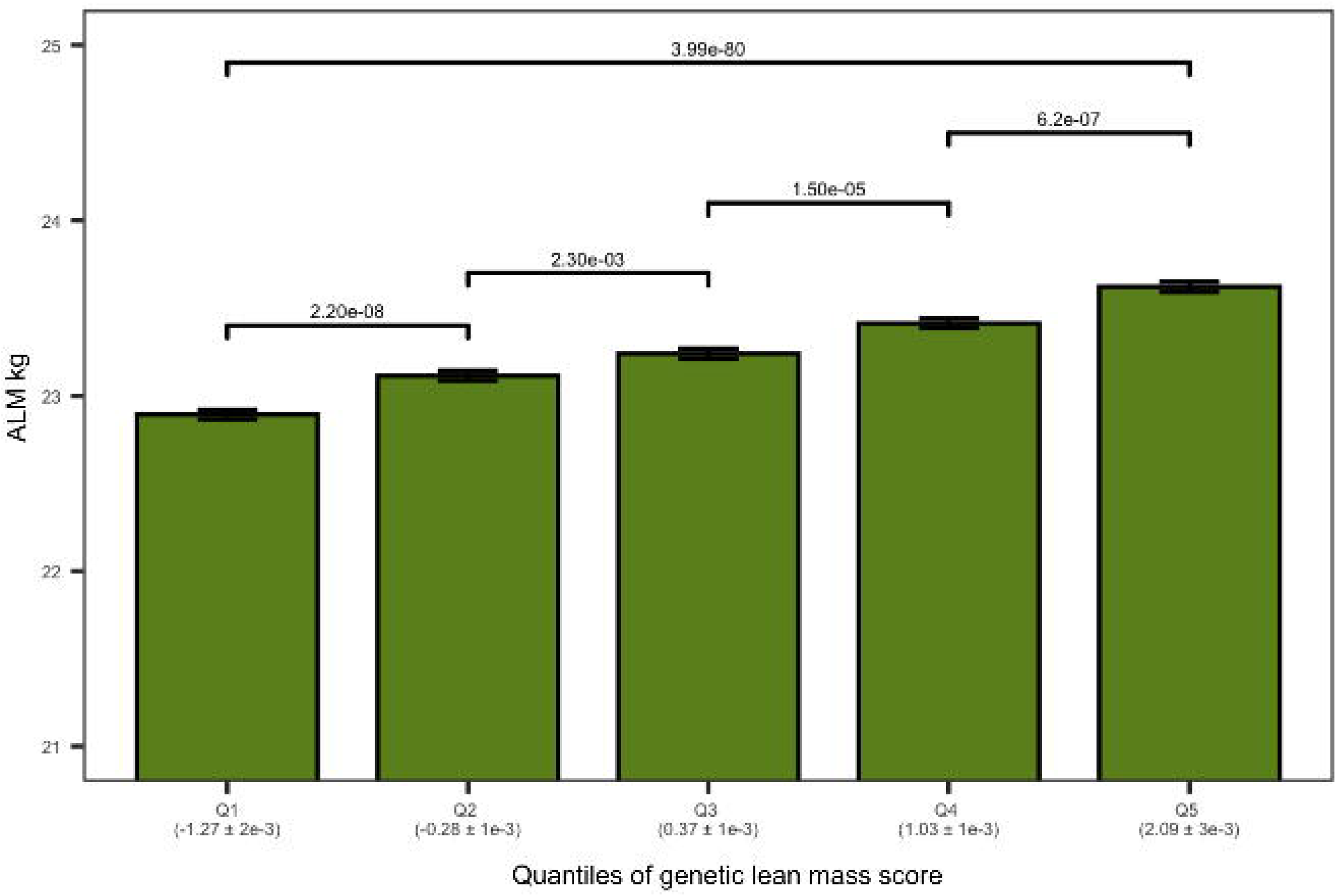
Genetic lean mass score affects the appendicular lean mass (ALM) in elderly humans. The plot shows the ALM (kg) of the elderly cohort on the vertical axis. The elderly cohort was ranked by genetic lean mass score and clustered in five quantiles (Q1 to Q5) (horizontal axis). The average genetic lean mass score (± standard error) of each quantile is shown in parenthesis below the horizontal axis. The overall quantile effect of the genetic lean mass score on ALM was tested with Kruskal-Wallis test and the resulting *P* value is presented on the top of horizontal line above the bars. The ALM median differences between the groups were tested using a Wilcoxon test; the significance level of each comparison is presented above the horizontal lines with a Holm adjusted *P* value.

We also asked if the variants identified in the middle-aged cohort were associated with ALM in the elderly. A GWAS in the elderly cohort replicated 5,291 variants based on their *P* value (*P* < 5 × 10^−8^) and allelic effect (*beta*); moreover, the replicated variants tagged 78% of the ALM loci of the middle-aged cohort (two tailed Fisher test *P* value < 2.2 × 10^−16^). Overall, the set of genomic loci in the elderly cohort appeared similar to that of the middle-aged adults, with the exception of an approximately 5 Mb region on chromosome 5 (Figure S4). This region showed a very strong association with the ALM variability in older adults (lowest *P* value = 4.50 × 10^−56^, *beta* = 0.12 ± 0.01 kg), and had a modest albeit significant association with the ALM of middle-aged individuals (lowest *P* value = 3.30 × 10^−11^) with an effect size of 0.07 ± 0.01 kg.

### 23 QTLs contribute to muscle weight variability in LG/J and SM/J strain-derived advanced intercross lines

We examined the weight of four hindlimb muscles of the LGSM AIL (F_34_ and F_50_-F_56_): tibialis anterior (TA), extensor digitorium longus (EDL), gastrocnemius and soleus. The LGSM AIL muscles showed extensive individual variability (Table 2); furthermore, the SNP heritabilities of the TA, EDL, gastrocnemius and soleus muscles were 0.39, 0.42, 0.31 and 0.30, respectively (Table 2). The genome mapping of LGSM AIL muscles yielded 23 QTLs (*P* < 6.45 × 10^−06^). The TA, EDL and gastrocnemius QTLs explained more than the 50% of the SNP heritability of each trait (Table S6). The soleus muscle phenotypic variability explained by QTLs was 23% of its SNP heritability. Three QTLs were shared between the four hindlimb muscles (chromosome 7, 11 and 13; (Figure 3); the QTL on chromosome 13 resulted in the strongest association (EDL *P* = 2.95 × 10^−21^), with its peak position at 104,435,003 bp, and the percentage of phenotypic variance explained by this locus was 5.2%; the SM/J allele conferred increased muscle mass (Figure 3). Furthermore, six QTLs were shared between two or three hindlimb muscles, while fourteen identified QTLs were only associated with one specific hindlimb muscle (Figure 3).

**Figure 3.**
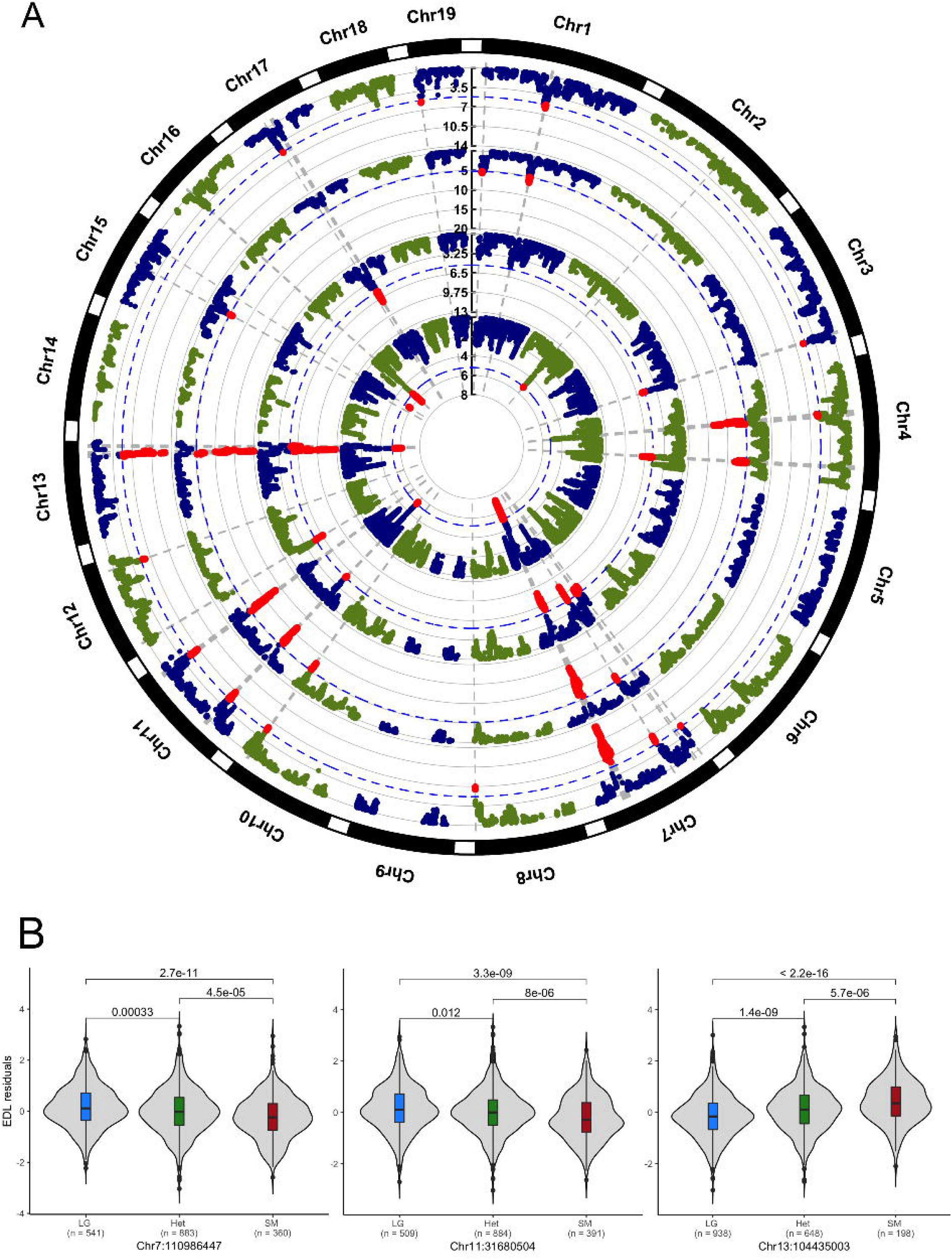
Muscle weight QTLs identified in mice of the LGSM AIL and density plot of the genotypes. The circle plot (A) shows from the outer to the inner ring the GWAS of the TA, EDL, gastrocnemius and soleus muscle weights. Chromosomal position of each SNP is shown in the outer black circle of the plot; chromosome names are shown outside as “Chr”. Dots within each chromosome space represent the association (–log10 *P value*) of each SNP tested. Dotted blue lines represent the genome-wide threshold (*P* < 6.45 × 10^−06^) of significance, and red dots above the genome-wide threshold are significantly associated SNPs. (B) Plots of the allelic effect of the *Skmw34*, *Skmw55* and *Skmw46* QTLs on the EDL muscle mass. These QTLs were identified for the four muscles investigated. Vertical axis represents the residual muscle mass adjusted for sex, age, dissector and long bone length of the hindlimb, and the horizontal axis shows the genotypes (LG/J homozygote, heterozygote and SM/J homozygote). Below the horizontal axis, the number of individuals with a given genotype is provided. The violin shapes within the plot area represent the distribution of individuals with the genotypes. Box whiskers represent minimum and maximum values, distance between a whisker and the top or bottom of the box contains 25% of the distribution, the box captures 50% of the distribution, and the bold horizontal line represents the median. Pairwise comparison *P* value (t-test) is shown above horizontal lines at the top of the plots.

**Table 2.**
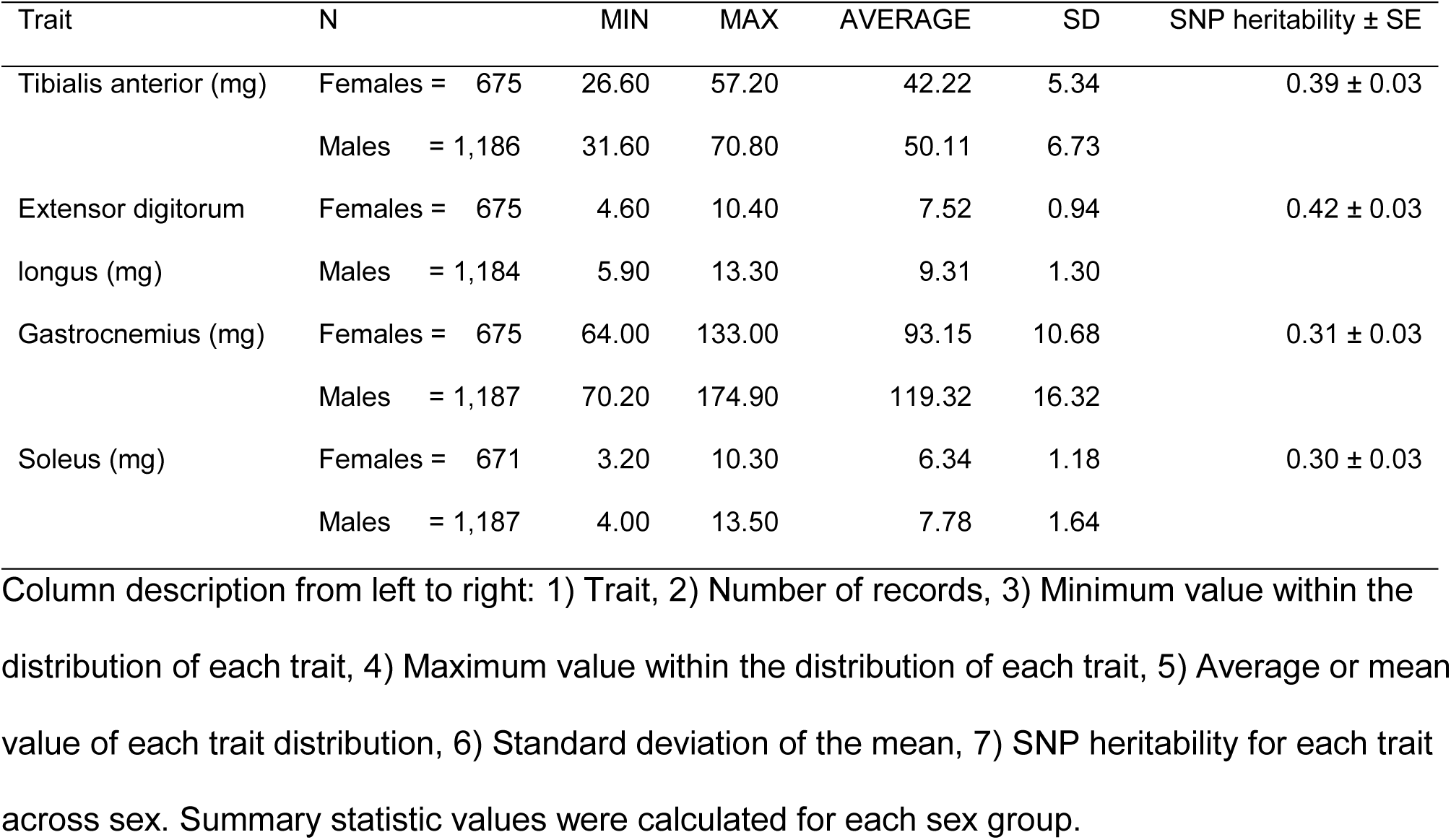
Summary of the LGSM AIL muscle traits

The mapping resolution was comparable to that attained in the previous study in the LGSM AIL cohort^59^. On average, mouse QTLs spanned 2.80 Mb (based on the 1.5 LOD interval) and encompassed 2,267 known genes (Table S7). The median number of genes per QTL was 55; more than half of the mouse QTLs contained a modest number of genes; however, 7 QTLs contained more than 100 genes each, and a single QTL located on chromosome 7 as many as 644 genes (Table S6). Although all mouse QTLs identified in the LGSM AIL contained SNPs, at least seven QTLs covered long genomic regions characterized as identical by descent between the LG/J and SM/J strains^58^. We also analyzed the LGSM AIL using a meta-analysis approach and identified 14 QTLs that on average were 3.78 Mb long. The majority of the QTLs from the meta-analysis, 12 out of the 14, overlapped with the findings of the mega-analysis (Figure S5). The meta-analysis results are shown in Table S8.

### Interspecies overlap between appendicular lean mass loci and muscle weight QTLs

The ALM mainly consists of the skeletal muscle of the extremities; however, other tissues also contribute. To test the hypothesis that ALM-associated genetic variants primarily affect skeletal muscle mass, we overlaid human ALM finding with the mouse where skeletal muscle was weighed directly. Specifically, we overlaid the captured genomic regions restricted by the significant SNPs used in the GWAS of each species. The mouse QTL regions were notably larger, partially due to the median distance between adjacent genetic markers of 126.9 Kb. Our analysis identified five syntenic regions associated with ALM in humans and hindlimb muscle mass in mice (Table 3). We used Fisher’s exact test to discover that the number of overlapping regions significantly (*P* = 0.0019; Table S9) exceeded that which could be expected by chance. This analysis permitted us to shorten the list of positional candidates. Assuming the same causative entity for an overlapping mouse and human locus, these five loci harbour 38 homologous genes (Table 3). Encouragingly, four of these five genomic loci tagged by rs148833559, rs9469775, rs4837613 and rs57153895 SNPs (Table S3) were replicated in the ALM of the elderly cohort (Table S10).

**Table 3.**
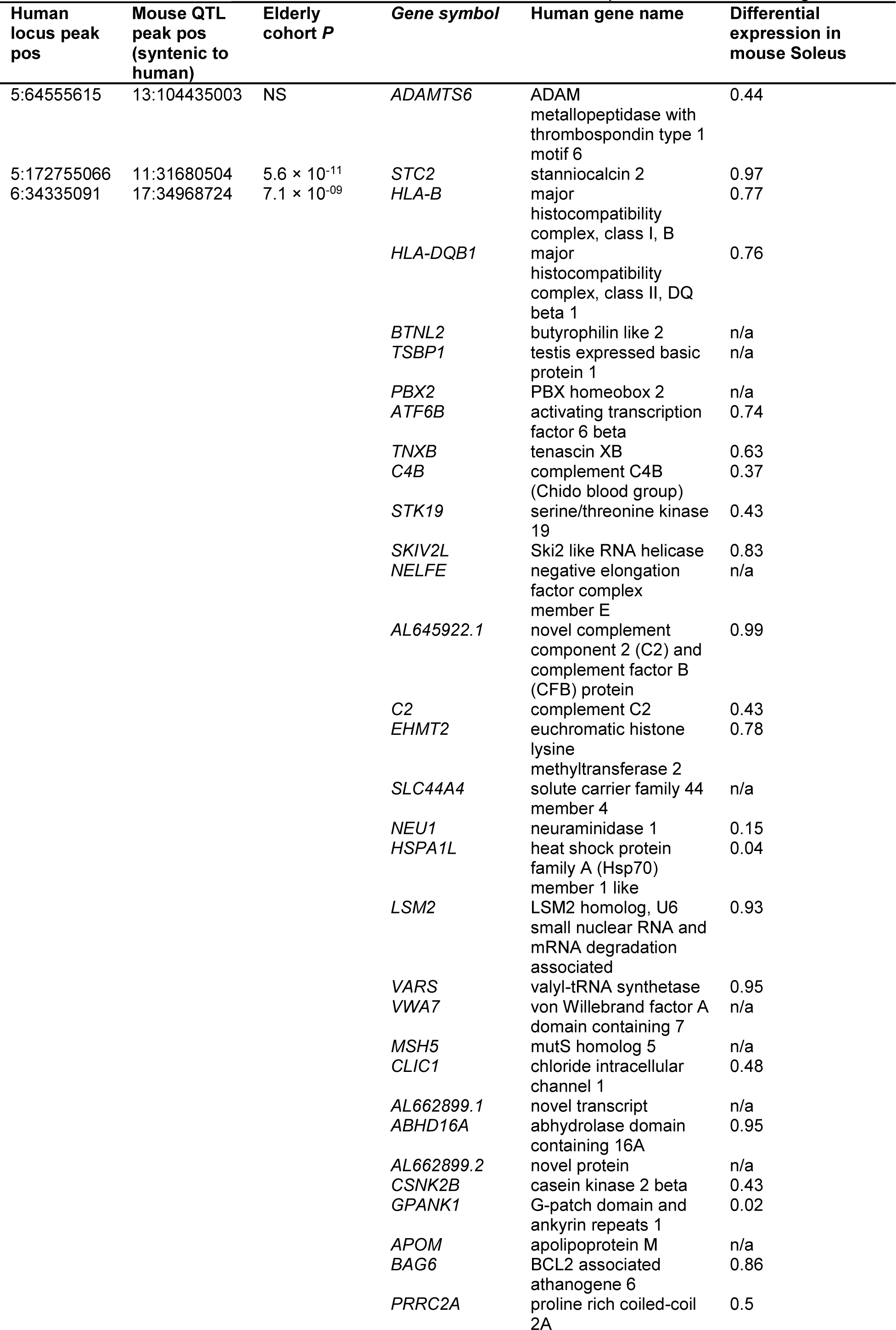
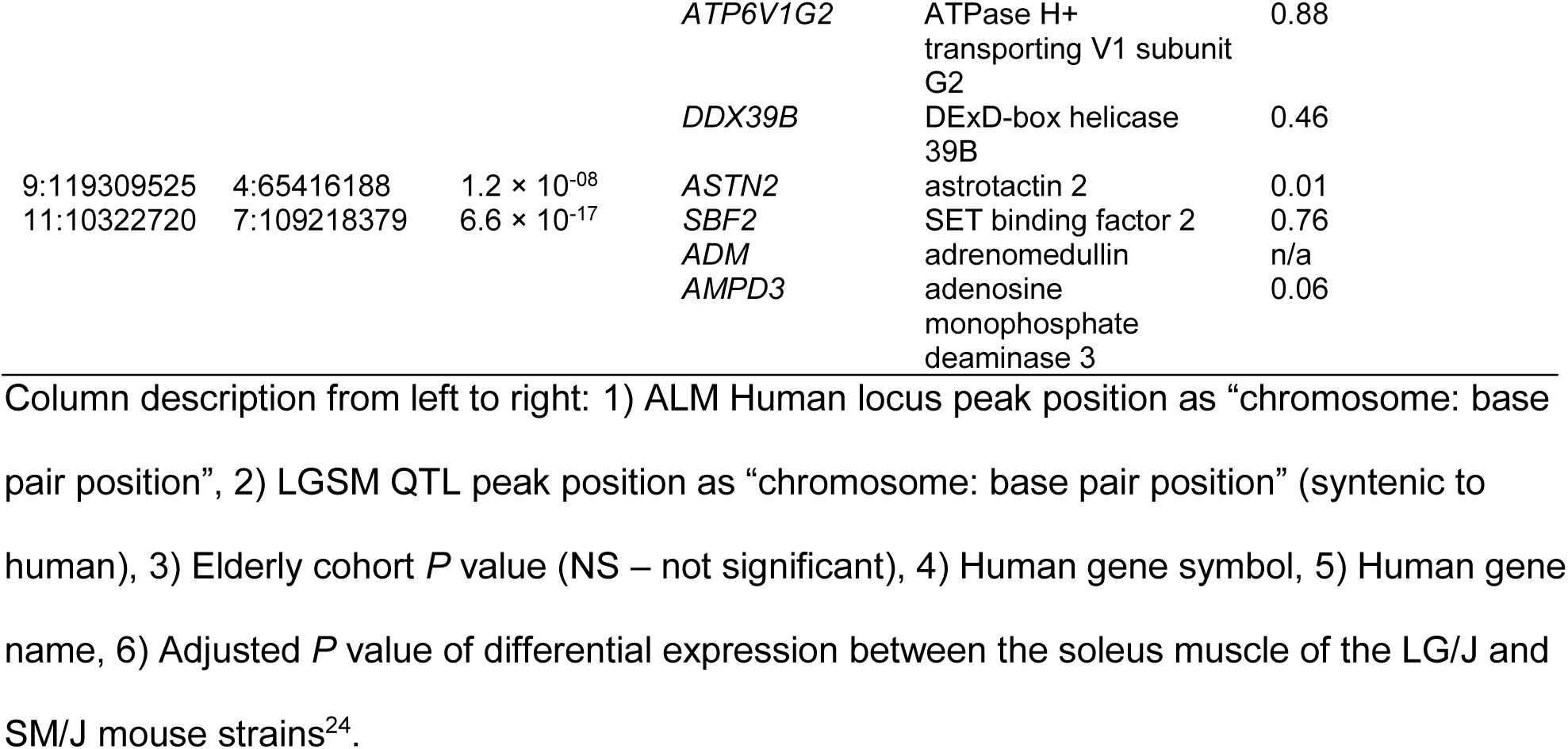
Syntenic regions between human and mouse QTLs and positional candidate genes

### Modifiers of *in vitro* myogenesis

We used siRNA-mediated gene knockdown in C2C12 cells to test if candidate genes affected myogenic differentiation. The *STC2*^85^, *CPNE1*^5^ and *SBF2*^86^ were prioritized for this assay because they were highlighted by both mouse and human GWAS. We assessed indices of myogenic differentiation (the number and length of the myotubes, and expression of myosin) of C2C12 cells. In total, 34,989 myotubes were identified and measured in 44 cell cultures (see Methods for details). The gene knockdown had a significant effect on myotube length, with *Cpne1 (P = 0.001, 95% confidence interval = 0.019-0.068, effect size = 0.024)* and *Stc2 (P = 0.015, 95% confidence interval= 0.007-0.066, effect size = 0.017)* showing an increase in length compared to the control cells (Figure 4). There was no significant difference for the *Sbf2* gene. The pattern of the effect on myosin expressing area was similar to that of myotube length but was not statistically significant (*P* = 0.21). The number of myotubes was also unaffected.

**Figure 4.**
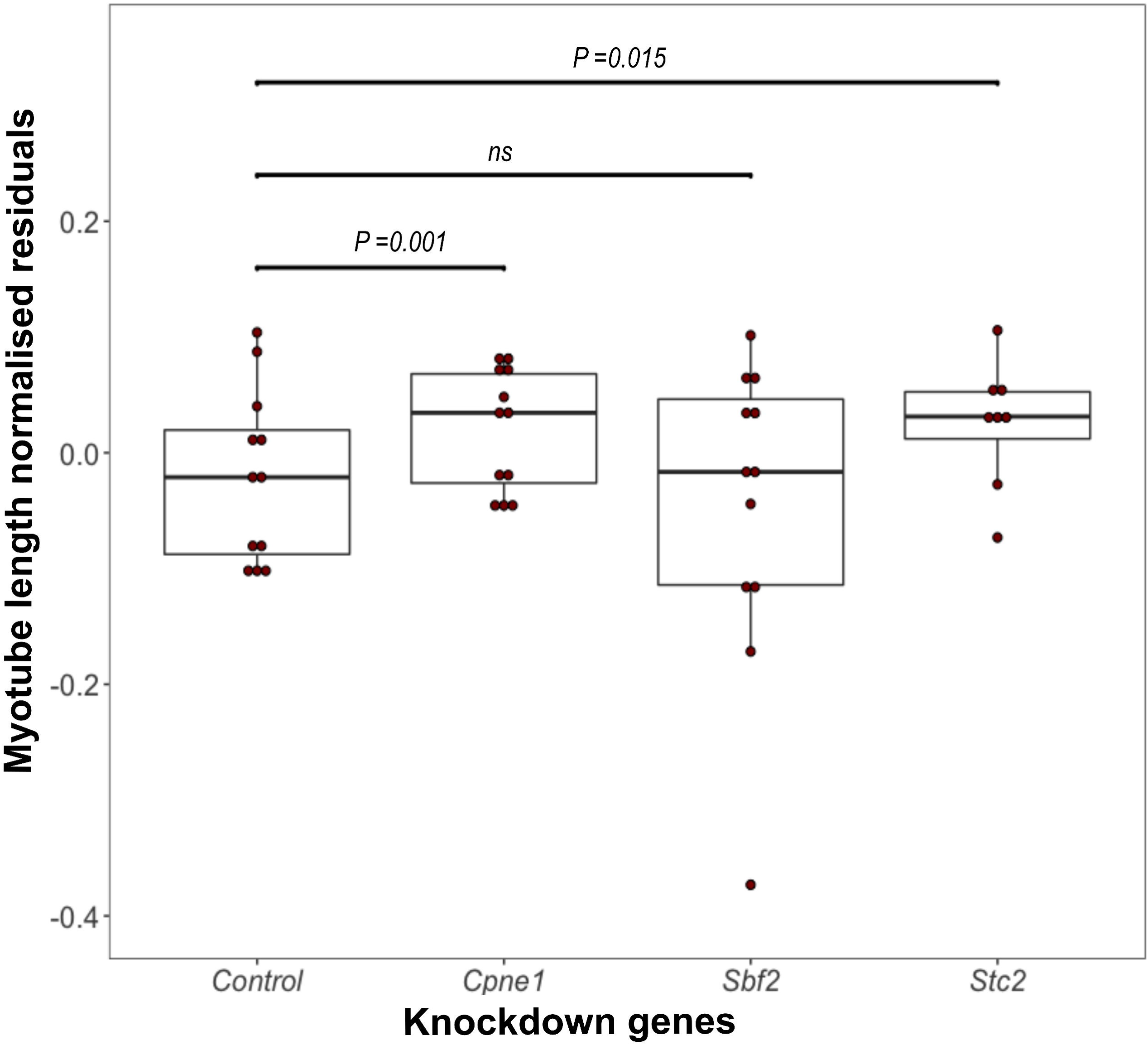
Gene knockdown effect on C2C12 myotube length. This figure shows the gene knockdown effect of the *Cpne1*, *Sbf2* and *Stc2* genes on myotube length. The overall effect of the gene knockdown on myotube length was tested using ANOVA and the resulting *P* value was 0.00017 (F3, 34985 =6.63). Vertical axis represents the myotube length (quantile normalised) residuals (adjusted for area analyzed and batch of cells), and the horizontal axis shows control and knockdown gene groups. Boxes represent the distribution of the myotube length for each group. Box whiskers represent minimum and maximum values within 1.5-fold interquartile range above the 75^th^ percentile and below the 25^th^ percentile; the box captures 50% of the distribution, and the bold horizontal line represents the median value of the myotube length normalized residuals distribution for each knockdown group. Each red dot represents a single cell culture sample for each knockdown group. Statistically significant t–test *P* values between control and knockdown genes are presented above horizontal lines. Effects without a statistically significant difference between the control and gene knockdown are presented as “ns”. *Cpne1* and *Stc2* knockdown groups were not different from each other (*P > 0.05*). *Sbf2* knockdown differed from *Cpne1 (P = 0.002)* and *Stc2 (P = 0.043)*.

## Discussion

The key findings of the present report are as follows: 1) we identified a set of over 180 loci associated with ALM, a substantial expansion in comparison to previous human studies. 2) There is a substantial overlap of the genetic effects between middle-aged and elderly subjects. 3) Integration of mouse and human GWAS indicates that skeletal muscle is the primary component affected by the ALM loci, facilitates prioritization of candidate genes, and helps prediction of their effect on cellular mechanisms underlying muscle mass variation. 4) *In vitro* studies validated two genes, *CPNE1* and *STC2*, as modifiers of muscle mass in humans.

We estimated SNP heritability for ALM to be of 0.36, which is lower than heritability estimates previously reported (0.44) ^8^, this difference could be due to different fixed effects used to estimate variance components. In total, we mapped 182 loci that collectively explain 24% of the SNP heritability of ALM. The most recent report, a meta-analysis of 47 independent cohorts (dbGAP), comparable in sample size but ranging in subjects aged 18 to 100 years, reported five significant associations with lean body mass^8^. Even fewer associations were detected in the earlier, small sample size studies^10; 12–14; 87^. However, our results indicate that ALM is a highly polygenic trait in humans. We hypothesize that multiple factors contributed to the improved locus detection in the present GWAS. We restricted subjects’ age to a narrow range, 38 to 49 years, minimizing the effects of the developmental and aging-related processes on phenotypic variance. The skeletal muscle is a dynamic tissue reaching its peak mass by late 20s, then a trend of decline emerges after 40s and accelerates about two decades later^1^. An estimated 30-50% decline in muscle mass can be expected between 40 and 80 years of age^88^. These developmental and aging-related changes are not linear in progression and therefore would hamper detection of loci even if accounted for in a linear model. In addition, unlike Zillikens and colleagues^8^, the data set we used was systematically collected as described by the UK Biobank project^15^ and we only employed bioelectric impedance measurements of lean mass. Furthermore, we used a LMM to test the effects of > 21 million variants (MAF > 0.001), and our analysis was adjusted for a different set of fixed effects than in previous research^8; 10; 12; 14^. Our analysis captured three loci identified by Zillikens and colleagues^8^, containing the *VCAM, ADAMTSL3* and *FTO* genes, suggesting that their effects are not influenced by age. We hypothesize that a combination of a homogeneous age group, the optimized genomic coverage and the method used to conduct this association analysis contributed to improved detection of loci in the present study.

The analyses presented here shed light into the complex genetic mechanisms behind the appendicular muscle mass of humans. In the past, concern was expressed about the reproducibility of association analyses of complex traits; however, an increasing number of human GWAS have shown that their findings are remarkably reproducible^89^. The present study provides further support for the reliability of association studies, demonstrating replication of 78% of ALM loci in the elderly cohort. Furthermore, we show that the genetic profile characterized by depletion of ALM-increasing alleles leads to a lower ALM in elderly individuals (Figure 2). Hence, it is conceivable that genetic architecture predisposing individuals to lower muscle mass may lead to elevated risk of sarcopenia^1^.

Combining two experimental models, mouse and human, facilitated prioritization of candidate genes for functional validation and indicated that skeletal muscle is the primary component of lean tissues affected by the identified loci. Furthermore, the mouse model revealed that genetic effects may not all be uniform across skeletal muscle tissue, instead some of the effects can be muscle type- or muscle-specific. To establish the association between the QTGs of the identified loci and the muscular phenotype, we focused on the overlapping human and mouse results. Integration of results from these two species permitted circumvention of the limitations imposed by the individual models. While human GWAS often identify loci containing single genes, it is often unclear which tissue is most relevant to the phenotype. Although mouse QTLs often contain multiple positional candidate genes, mice can be used as experimental models to identify loci specifically associated with skeletal muscle. In this study, we used a mouse model to show that the association with hindlimb skeletal muscle mass was specifically related to differences in the cross-sectional area of the constituent muscle fibers, rather than to the number of fibers in the muscle. This is because between the two founders of the LGSM AIL, the LG/J strain compared to the SM/J strain shows over 50% larger cross-sectional area of muscle fibers, but no difference in the number of fibers in soleus muscle^21^. Hence, it is conceivable that the QTGs of the majority of the overlapping loci affected hindlimb muscle mass specifically via the hypertrophy of muscle fibers. Such prioritization between the two cellular mechanisms of muscle mass variability is important because genes specifically influencing cross-sectional area of muscle fibers can be targeted pharmacologically to prevent and reverse atrophy of muscle fibers in aging muscle^90^. In humans, the bone, muscle and skin tissues contribute to lean mass determined by bioelectric impedance. Approximately 1-2 mm thick skin^91^ constitutes a rather minor component of lean mass compared to the size of human extremities. The long bones determine axial dimensions of a limb but we adjusted for that to minimize bone effect on variation of lean mass. Whereas the magnetic resonance imaging (MRI) assessed muscle mass accounts for ∼38% of body weight in humans, and the MRI data strongly and positively correlates with the estimates of bioelectric impedance^92^. This, collectively with the overlap of the ALM loci and mouse muscle QTLs, provides a strong support for the notion that skeletal muscle is the primary tissue affected by the ALM loci.

For the functional validation we prioritized three candidate genes (*STC2*, *SBF2* and *CPNE1*) implicated by both human and mouse analyses. *STC2* had the largest effect size on the ALM (beta = 0.88 ± 0.13 kg; Table S3) with the minor allele (A) of a missense SNP (rs148833559 (A/C) associated with the increase in ALM. Prediction tools (SIFT^53^, PolyPhen^51^, CADD^93^, and REVEL^94^) suggested a detrimental consequence of rs148833559 on STC2 structure. *SBF2* has been linked to Charcot-Marie-Tooth hereditary motor and sensory neuropathy^86^, and is expressed in skeletal muscle and associated with a cis-eQTL^55^. Although little is known about *CPNE1,* it is an intriguing candidate because the minor allele of the missense variant (rs12481228) is predicted by SIFT^53^ and PolyPhen^52^ to be detrimental on the structure of CPNE1. That allele was associated with increased ALM in the middle-aged cohort, and a frameshift variant (rs147019139) leading to premature stop codon was also associated with an increase in ALM in the elderly cohort (Table S10). Furthermore, in a previous GWAS using outbred (CFW) mice^5^, *Cpne1* was implicated in hindlimb muscle mass. To validate these QTGs for their effects on skeletal muscle, we tested the siRNA-mediated knockdown effect on myogenesis *in vitro*. A knockdown of two genes, *CPNE1* and *STC2*, increased the length of the myotubes. Although it is not completely understood how changes in the indices of *in vitro* myogenesis correlate with the fiber hypertrophy and/or hyperplasia *in vivo*, our findings implicate an upregulation of myogenic differentiation. We interpret this *in vitro* observation as being consistent with the allelic effect of the two loci identified in human GWAS. *CPNE1* encodes for Copine 1, a soluble calcium-dependent membrane-binding protein^95^ expressed in skeletal muscle^24^. *STC2* encodes Stanniocalcin 2, a homodimeric glycoprotein hormone abundantly expressed in skeletal^55^ and cardiac muscle^96^, and involved in regulation of IGF1 through interaction with pregnancy-associated plasma protein-A^97^. A suppressive role of *STC2* is consistent with reduced muscle mass in the STC2 overexpressing mice^85^. Hence, our analyses and recent reports provide support for *CPNE1* and *STC2* as suppressors of muscle mass development and/or maintenance in humans.

In conclusion, the present study integrated human and mouse GWAS and used *in vitro* validation to further interrogate a subset of the genes implicated in both species. Our results revealed over 180 genomic loci contributing to ALM in middle-aged humans. The effects of the majority of these loci persist in the elderly human population. Integration of human and mouse data also highlighted candidate genes affecting skeletal muscle mass in mammals. Two genes, *CPNE1* and *STC2* were confirmed to be modifiers of *in vitro* myogenesis.

## Supplemental Data

Supplemental Data include 5 figures, 10 tables and 1 macro script.

## Acknowledgements

The authors would like to acknowledge Dr David A. Blizard for his role in the development of the ideas that led to this study and feedback on the manuscript, Professor Helen Macdonald for valuable advice on study design, Dr Leslie R. Noble for help with the UK Biobank data, and Dr Joseph P. Gyekis for help genotyping cohort 2 mice. The authors would like to acknowledge funding from the University of Aberdeen for the Maxwell computer cluster, the Elphinstone and IMS studentship for AIHC; a Schweppe Foundation Career Development Award (AAP), and the NIH (NIAMS (AL: R01AR056280) and NIDA (AAP:R01DA021336, AAP:R21DA024845, AAP:T32MH020065, NMG:F31DA03635803), NIGMS (NMG:T32GM007197), NHGRI (MA:R01 HG002899)).

## Web Resources

Functional Mapping and Annotation of Genome-Wide Association Studies (FUMA GWAS)^46^. URL: http://fuma.ctglab.nl/ Ensembl^54^. URL: https://www.ensembl.org Gene Tissue Expression Project (GTEx) portal^36^. URL: https://gtexportal.org/home/

