## Supplemental Materials and Methods for "Genome-wide associations reveal human-mouse genetic convergence and modifiers of myogenesis, *CPNE1* and *STC2*"

**Macro script– Myotube length analysis (Fiji)**

//get the title of the image & put into a variable

imName=getTitle();

//First step - open the channel with myotube staining & make the largest possible rectangular ROI comprising "good" image data

// Avoid artefacts, peeling-away cells and blurred or bleached areas.

// Duplicate this image & save and run the macro

run("RGB to Luminance");

//Bandpass FFT to get rid of vertical & horizontal lines

run("Bandpass Filter...", "filter_large=80000 filter_small=6 suppress=Vertical tolerance=3");

run("Bandpass Filter...", "filter_large=80000 filter_small=6 suppress=Horizontal tolerance=3");

//Rolling ball background subtraction

run("Subtract Background...", "rolling=200");

// Threshold the image - added a step for the user to alter manually if needed

setAutoThreshold("Li dark");

waitForUser("is the threshold OK?");

setOption("BlackBackground", false);

run("Convert to Mask");

//Measure the % area

run("Set Measurements...", "area mean min area_fraction display redirect=None decimal=3");

run("Select All");

run("Measure");

// Save results in variables for later

roiArea=getResult("Area");

areaFraction=getResult("%Area");

//Skeletonization

run("Skeletonize (2D/3D)");

waitForUser("click if skeletonize finished");

run("Analyze Skeleton (2D/3D)", "prune=none calculate show");

selectWindow("Results");

// Add key results to a summary table (or make it if there isn't one already open)

title1 = "SummaryData";

title2 = "["+title1+"]";

f = title2;

if (isOpen(title1)){

// do nothing for now

}

else {

//If the table doesn't exist yet, make a new one to collate summary data.

// Get the image name, Area of the ROI and the rest of the summary results

run("Table...", "name="+title2+" width=400 height=300");

print(f, "\\Headings:Image\tROIArea\tPercent_Green");

// print(f, "\\Headings:Image\ROIArea\tPercent_Green\tNumPaths\tNumPaths>20\tLongestShortestPath\tLongestShortest>20");

}

// Print summary of results to a table

print(f,imName +"\t"+ roiArea +"\t"+areaFraction);
