## Supplemental Materials for "Genome-wide associations reveal human-mouse genetic convergence and modifiers of myogenesis, *CPNE1* and *STC2*"

**Supplemental Data**

- Figure S1: QQ plot -ALM associations *P* values (middle-aged cohort)
- Figure S2: Myotube length analysis
- Figure S3: Negative controls of the genetic lean mass scores tested in an elderly cohort from the UK Biobank
- Figure S4: Map of genome associations with the appendicular lean mass (ALM) in elderly humans
- Figure S5. LGSM AIL genome mapping by mega and meta-analysis
- Table S1. Summary statistics for the elderly human cohort
- Table S2. Quantitative PCR primers
- Table S5. Summary of the negative controls for the genetic lean mass

Table S9. Contingency table of syntenic regions (Fisher test)

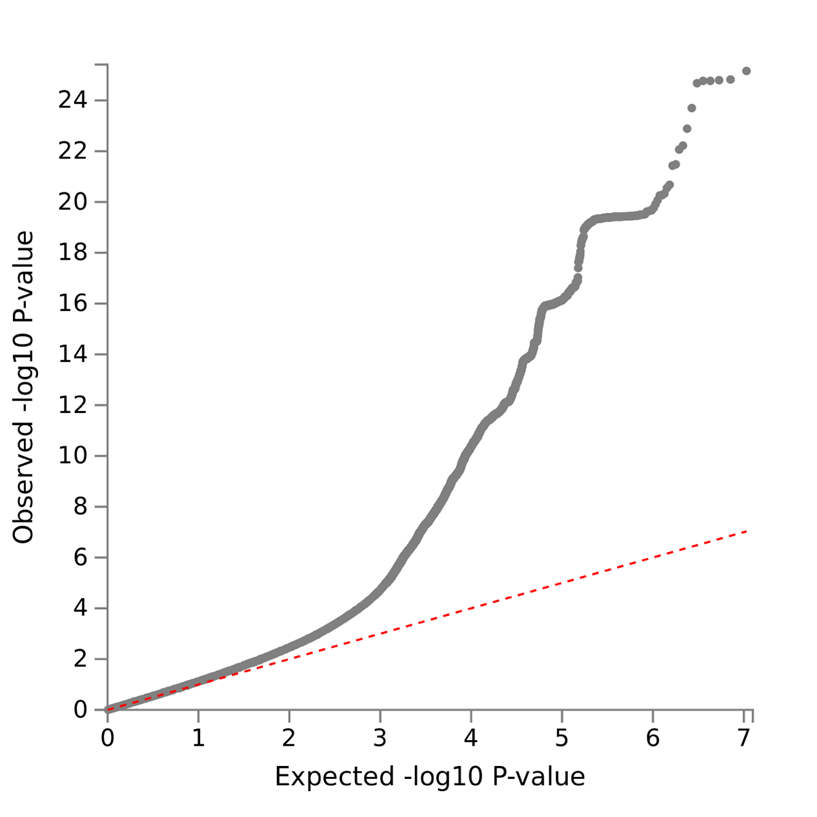

Figure S1. QQ plot of the GWAS on the appendicular lean mas of 85,750 individuals from the UK Biobank project. Lambda (λ_GC_) = 1.20.

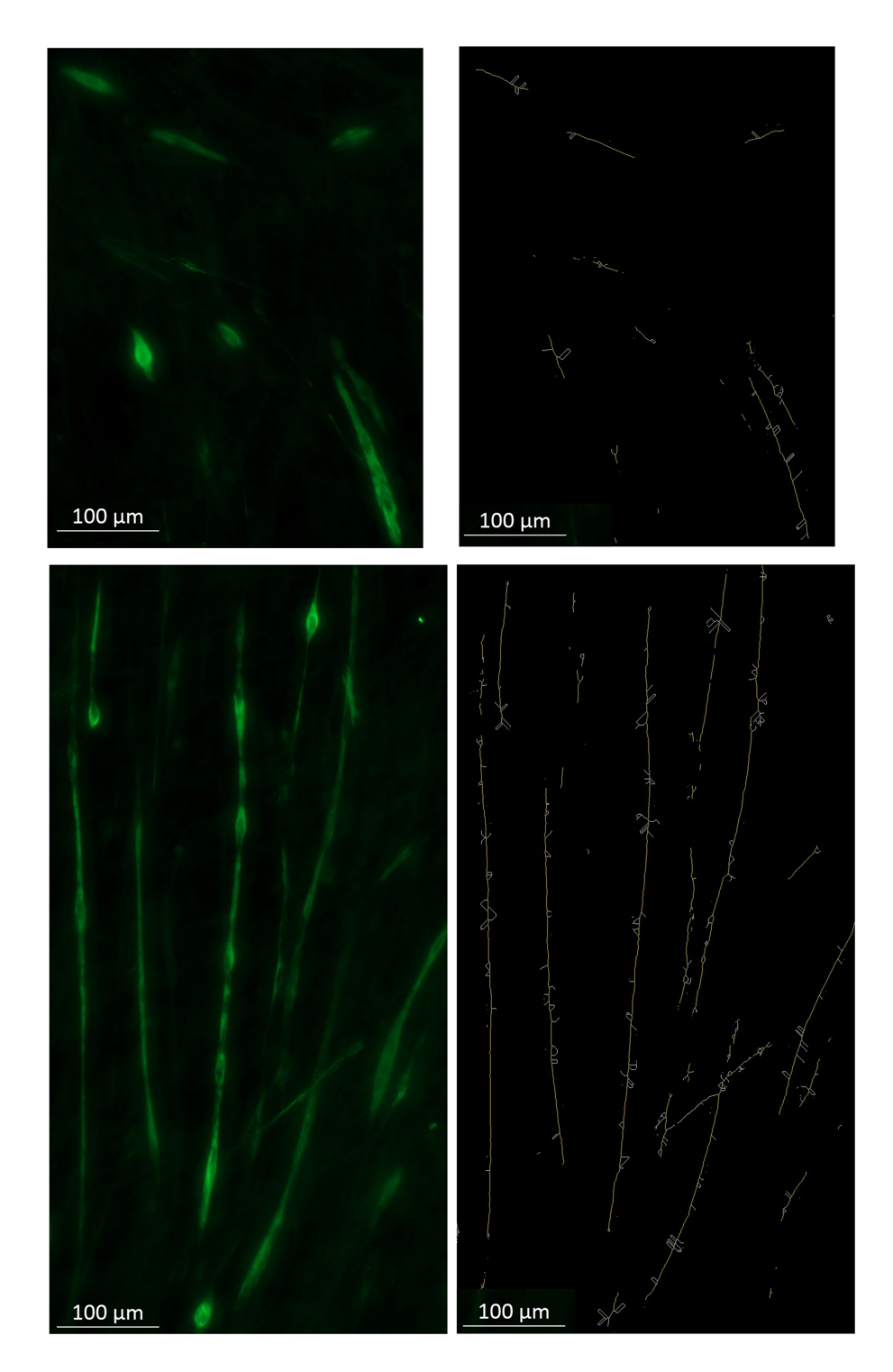

Figure S2. Myotube length analysis.

Figure shows examples of two contrasting regions (top and bottom) of myosin stained C2C12 cells. The original image presented on the left and the result of the image analysis on the right. The longest shortest path is the longest distance between two points on any skeleton using the shortest route possible. This has previously been applied to images of plant roots with an R^2^ value of over 0.9 between manual and automatic measurements of length^1^. The resulting skeletons are displayed in 2 colours, with the longest shortest path^1^ (the measure of myotube length) highlighted in yellow and the branches of the skeleton (white) not included in the measure shown in white. To perform the analysis, first a region of interest with no artefacts (other than translucent out-of-focus areas) is selected. The macro was then applied. The first step of the macro performed a rolling-ball background subtraction to ensure each myotube stands out clearly from any background fluorescence. An automatic threshold (Li's Minimum Cross Entropy^2^) was then used to segment the green areas of the myotubes. The area was measured and calculated as a percentage.

Following this the Analyze Skeleton plugin^3^ identified the central axis of each region and the resulting “skeletons” were analysed to measure the length using the longest shortest path. Results were filtered by length to remove myocytes and noise, leaving only the skeletons long enough to be mytotubes for the analysis.

Whilst the myotubes in this experiment are complex and likely to overlap or have small gaps in segmentation this approach provided an unbiased quantitative estimate of the length and number of myotubes in each experiment.

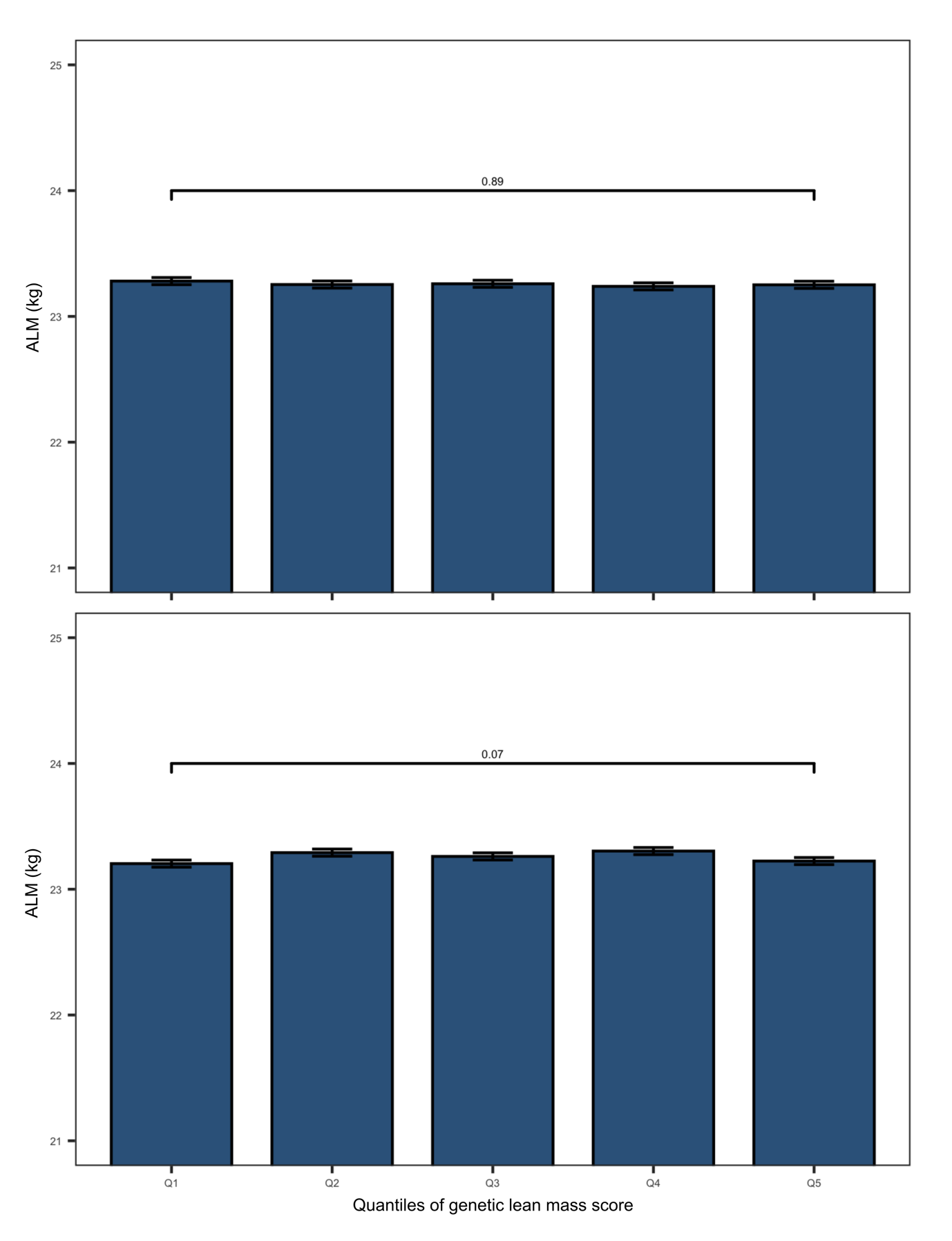

Figure S3. Negative controls of the genetic lean mass scores tested in an elderly cohort from the UK Biobank. The figure shows two negative controls. Each bar plot shows the ALM (kg) of the elderly cohort in the vertical axis. The elderly cohort was ranked based on the genetic lean mass scores and clustered in five quantiles (Q1 to Q5) (horizontal axis). The overall quantile effect of the genetic lean mass score on appendicular lean mass was tested with Kruskal-Wallis test and the resulting *P* value is presented on the top horizontal line above the bars.

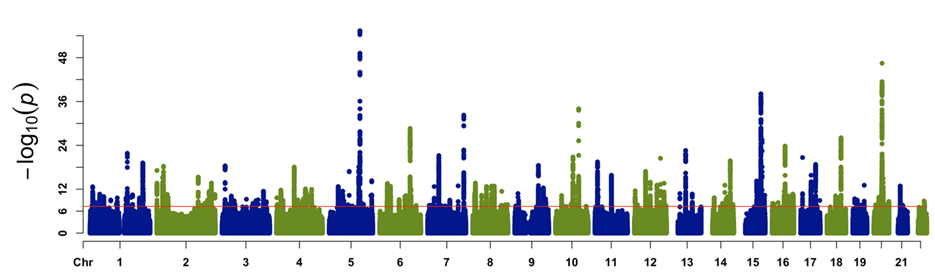

Figure S4. Map of genome associations with the appendicular lean mass (ALM) in elderly human cohort from the UK Biobank. Significance level is presented on the vertical axis, while the chromosomal position of each genetic marker is shown on the horizontal axis. Red line across the plot represents the genome wide threshold of significance (*P* < 5 x 10^-8^). This plot shows the association of variants with MAF > 0.001.

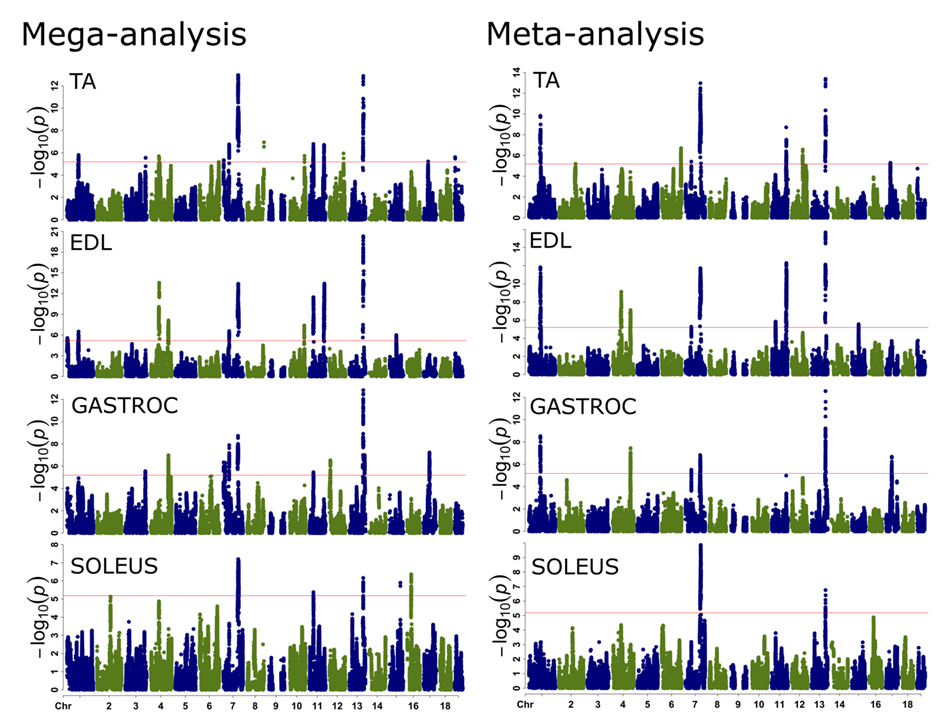

Figure S5. Muscle weight QTLs identified in mice of the LGSM AIL. Left panels show rectangular Manhattan plots that resulted from the mega-analysis of the four LGSM AIL muscles (TA= tibialis anterior, EDL= extensor digirotum longus, GASTROC: gastrocnemius). Right panels show rectangular Manhattan plots that resulted from the meta-analysis of the four LGSM AIL muscles. Significance level is presented on the vertical axis, while the chromosomal position of each genetic marker is shown on the horizontal axis. Red line across the plot represents the genome wide threshold of significance (*P* < 6.45 x 10^-06^).

Table S1. Summary statistics for the elderly human cohort

| Trait | N | | MIN | MAX | AVERAGE | SD |
| --- | --- | --- | --- | --- | --- | --- |
| Age (years) | Females = | 94,229 | 60.00 | 71.08 | 64.48 | 2.84 |
|  | Males = | 87,633 | 60.00 | 73.67 | 64.67 | 2.85 |
| ALM (kg) | Females = | 92,536 | 11.10 | 57.00 | 19.13 | 2.41 |
|  | Males = | 85,563 | 14.20 | 58.50 | 27.72 | 3.82 |
| Arm lean mass (kg) | Females = | 92,548 | 1.00 | 26.00 | 2.25 | 0.30 |
|  | Males = | 85,578 | 1.20 | 7.00 | 3.52 | 0.53 |
| Leg lean mass (kg) | Females = | 92,561 | 2.30 | 16.10 | 7.34 | 0.94 |
|  | Males = | 85,589 | 5.50 | 23.80 | 10.36 | 1.43 |
| WBF (kg) | Females = | 92,546 | 5.00 | 94.70 | 27.34 | 9.30 |
|  | Males = | 85,403 | 5.00 | 108.10 | 22.88 | 7.97 |
| Leg (cm) | Females = | 93,944 | 44.00 | 135.00 | 75.77 | 4.24 |
|  | Males = | 87,306 | 35.00 | 110.00 | 82.89 | 4.52 |

Column description from left to right: 1) Trait, 2) Number of records, 3) Minimum value within the distribution of each trait, 4) Maximum value within the distribution of each trait, 5) Average or mean value of each trait distribution, 6) Standard deviation. Summary statistic values were calculated for each sex group.

Table S2. Quantitative PCR primers

| Gene | Forward | Reverse |
| --- | --- | --- |
| *Cpne1* | 5’-GGACTGAACGTGTTCGCAAC-3’ | 5’-ACACGGCTGTCCTTTAGCTC-3’ |
| *Sbf2* | 5’-AGCCTGGTGTTGGTATCCAG-3’ | 5’-GTCTCCTGCACCCAAGGAAA-3’ |
| *Stc2* | 5’-TGACCCTGGCTTTGGTGTTT-3’ | 5’-GACTTTCCCTGGGCATCGAA-3’ |
| *Actb* | 5’-GGTGGGAATGGGTCAGAAGG-3’ | 5’-GTACATGGCTGGGGTGTTGA -3’ |

Table S5. Negative control for the genetic lean mass score tested on elderly adults.

| Replicate | df | Χ2 | P value | Quantile (genetic lean mass score ± se) | N | AVERAGE | SD | SE |
| --- | --- | --- | --- | --- | --- | --- | --- | --- |
| Control 1 | 4 | 1.11 | 0.89 | Q1 (-0.18 ± 3.90e-05) | 35,652 | 23.28 | 5.35 | 0.03 |
|  |  |  |  | Q2 (-0.17 ± 6.40e-06) | 35,561 | 23.25 | 5.35 | 0.03 |
|  |  |  |  | Q3 (-0.17 ± 5.29e-06) | 35,629 | 23.26 | 5.31 | 0.03 |
|  |  |  |  | Q4 (-0.16 ± 6.46e-06) | 35,579 | 23.24 | 5.31 | 0.03 |
|  |  |  |  | Q5 (-0.15 ± 4.96e-05) | 35,678 | 23.25 | 5.35 | 0.03 |
| Control 2 | 4 | 5.16 | 0.27 | Q1 (0.56 ± 4.62e-05) | 35,575 | 23.21 | 5.32 | 0.03 |
|  |  |  |  | Q2 (0.57 ± 7.64e-06) | 35,616 | 23.26 | 5.35 | 0.03 |
|  |  |  |  | Q3 (0.58 ± 6.66e-06) | 35,606 | 23.29 | 5.34 | 0.03 |
|  |  |  |  | Q4 (0.58 ± 8.28e-06) | 35,674 | 23.23 | 5.30 | 0.03 |
|  |  |  |  | Q5 (0.59 ± 3.67e-05) | 35,628 | 23.29 | 5.35 | 0.03 |
| Control 3 | 4 | 3.23 | 0.52 | Q1 (-0.03 ± 3.84e-05) | 35,639 | 23.25 | 5.32 | 0.03 |
|  |  |  |  | Q2 (-0.02 ± 7.26e-06) | 35,613 | 23.21 | 5.29 | 0.03 |
|  |  |  |  | Q3 (-0.02 ± 6.17e-06) | 35,623 | 23.29 | 5.35 | 0.03 |
|  |  |  |  | Q4 (-0.01 ± 7.34e-06) | 35,603 | 23.28 | 5.38 | 0.03 |
|  |  |  |  | Q5 (0.00 ± 4.99e-05) | 35,621 | 23.26 | 5.33 | 0.03 |
| Control 4 | 4 | 8.80 | 0.07 | Q1 (0.03 ± 4.56e-05) | 35,622 | 23.20 | 5.33 | 0.03 |
|  |  |  |  | Q2 (0.04 ± 7.39e-06) | 35,600 | 23.29 | 5.35 | 0.03 |
|  |  |  |  | Q3 (0.04 ± 5.99e-06) | 35,629 | 23.26 | 5.34 | 0.03 |
|  |  |  |  | Q4 (0.05 ± 7.22e-06) | 35,642 | 23.30 | 5.34 | 0.03 |
|  |  |  |  | Q5 (0.06 ± 5.09e-05) | 35,606 | 23.22 | 5.30 | 0.03 |
| Control 5 | 4 | 3.11 | 0.54 | Q1 (-0.05 ± 2.54e-04) | 35,630 | 23.27 | 5.34 | 0.03 |
|  |  |  |  | Q2 (0.03 ± 6.39e-05) | 35,623 | 23.26 | 5.31 | 0.03 |
|  |  |  |  | Q3 (0.07 ± 5.49e-05) | 35,595 | 23.28 | 5.36 | 0.03 |
|  |  |  |  | Q4 (0.11 ± 6.69e-05) | 35,632 | 23.26 | 5.33 | 0.03 |
|  |  |  |  | Q5 (0.18 ± 2.32e04) | 35,619 | 23.21 | 5.32 | 0.03 |

Column description from left to right: 1) Replicates 2) Degrees of freedom for Kruskal-Wallis test, 3) Χ^2^ for Kruskal-Wallis, 4) *P* value for Kruskal-Wallis, 5) Five groups based on the ascending ranking genetic lean mass scores for each replicate, 6) Number of samples, 7) Average appendicular lean mass, 8) Standard deviation, 9) Standard error.

Table S9. Contingency table of syntenic regions

|  | Number of overlapping loci | Number of non-overlapping loci |
| --- | --- | --- |
| Mouse loci | 5 | 18 |
| Human loci | 5 | 177 |

Columns show the observed number of overlapping and not overlapping loci between muscle QTLs in mice and ALM loci in humans (rows). These counts were used for a Fisher test to determine if overlapping number of loci was consequence of randomness. Null hypothesis was rejected at *P* = 0.0019; odds ratio: 9.62, and 95%CI for odds ratio: 2.54 - 36.85.
